# Thermal cycling-hyperthermia ameliorates cognitive impairment of intracerebroventricular Aβ_25-35_-induced Alzheimer’s disease in C57BL/6 mice

**DOI:** 10.1101/2022.07.22.500895

**Authors:** Yu-Yi Kuo, Wei-Ting Chen, Guan-Bo Lin, You-Ming Chen, Hsu-Hsiang Liu, Chih-Yu Chao

## Abstract

Despite continuation of some controversies, Alzheimer’s disease (AD), the most common cause of dementia nowadays, has been widely believed to derive mainly from excessive β-amyloid (Aβ) aggregation, that would increase reactive oxygen species (ROS) and induce neuroinflammation, leading to neuron loss and cognitive impairment. Existing drugs on Aβ have been ineffective or offer only temporary relief at best, due to blood-brain barrier or severe side effects. The study employed thermal cycling-hyperthermia (TC-HT) as an alternative AD therapy and compared its effect with continuous hyperthermia (HT) *in vivo*. It established an AD mice model via intracerebroventricular (i.c.v.) injection of Aβ_25-35_, proving that TC-HT is much more effective in alleviating its performance decline in Y-maze and NOR test, in comparison with HT. In addition, TC-HT also exhibits a better performance in decreasing the hippocampal Aβ and BACE1 expressions as well as the neuroinflammation markers Iba-1 and GFAP levels. Furthermore, the study finds that TC-HT can elevate more protein expressions of IDE and antioxidative enzyme SOD2 than HT. Besides, after establishment of neuroprotective mechanism, removal of TC-HT-induced ROS can further augment protection of neural cells against Aβ. In sum, the study proves the potential of TC-HT in AD treatment, which can be put into clinical application with the use of focused ultrasound (FUS).

## Introduction

Alzheimer’s disease (AD) has been the most common cause of dementia nowadays, accounting for over 60% of all the cases [1]. One of the most notable hallmarks of AD pathogenesis is the accumulation in brain of toxic β-amyloid (Aβ) peptides [2], which are cleaved from the amyloid precursor protein (APP) by β- and γ-secretase [3]. Such abnormal Aβ deposits can activate neuroinflammation [4], generate reactive oxygen species (ROS) [5, 6], and cause neuron dysfunction and synapse loss, leading to memory loss and cognitive impairment [7, 8]. Therefore, Aβ appears to be the foremost target in AD therapy [9].

Currently, drug therapy is the mainstream AD treatment, aiming to suppress the generation [10], disrupt the aggregation [11], and promote the clearance [12] of Aβ. Drugs on proteases, such as insulin degrading enzyme (IDE), for alleviating Aβ degradation also have significant potential in AD treatment [13]. However, at present AD drugs have high failure rates, due to difficulty penetrating blood-brain barrier, on top of severe side effects, such as liver toxicity, strong autoimmune response, or carcinogenicity [14]. Hence, there is a need for a safe and effective alternative AD therapy.

Heat has long been regarded as an alternative therapy with significant potential, given its proven therapeutic effect in physiotherapy, ophthalmology, and cardiology [15]. Hyperthermia (HT), for instance, has been applied effectively in cancer treatment, either for directly inhibiting tumor growth [16] or supplementing radiotherapy and certain chemotherapies to augment their efficacy [17]. In addition, moderate thermal stress can stimulate the expression of neuroprotective proteins. For example, HT is capable of increasing the expression of heat shock protein (HSP) family [18], some of whose members can protect neuron cells against various stresses to achieve neuroprotection [19, 20]. Besides, HT treatment is conducive to increase the levels of IDE [21] and antioxidant proteins, such as nuclear factor erythroid 2–related factor 2 (Nrf2) and superoxide dismutase 2 (SOD2) [22, 23].

However, continuous high-temperature exposure can be thermotoxic to human neural cells, according to some reports [24]. Heat stroke, featuring abnormally high core body temperature (> 40°C), may cause cerebellar injury, hypertension, and central nervous system derangement [25, 26]. Whole-body HT treatment has also been reported to cause neurological and systemic complications, such as nausea, impaired pressure-flow autoregulation, brain edema, and arrhythmias [27]. On the other hand, although a study showed that long-term heat exposure could reduce the risk of memory loss among middle-aged men, it was time-consuming without certain curative effect [28]. Therefore, it is imperative to find a tolerable and effective thermal treatment for AD.

Our previous study put forth a novel thermal treatment, thermal-cycling hyperthermia (TC-HT), demonstrating its therapeutic effect in anticancer treatment [29, 30] and neuroprotection [31] *in vitro*, with minimal side effects. The study applied TC-HT to treat mice with AD, induced by intracerebroventricular (i.c.v.) injection of Aβ_25-35_, for comparison with the therapeutic effect of HT. In both cases, heat was applied to mouse brain specifically, to avoid potential side effect of whole-body HT.

In this paper, behavioral tests show that TC-HT significantly improves Aβ-induced memory impairment, with an effect much better than HT. In addition, TC-HT also exhibits a much better performance than HT in decreasing Aβ and β-secretase (BACE1) expressions, suppressing neuroinflammation markers ionized calcium-binding adapter molecule 1 (Iba-1) and glial fibrillary acidic protein (GFAP) levels, and increasing the expression levels of IDE and SOD2 enzymes. Furthermore, the study shows that removal of TC-HT-induced ROS can further augment the protective effect of TC-HT in an *in vitro* neural model. In sum, these results underscore the potential of TC-HT as an alternative AD therapy.

## Materials and Methods

### Ethics statement

All animal experiments were conducted according to the regulations approved by the Animal Ethical Committee of National Taiwan University, and the approval protocol number was NTU-109-EL-174. Mice were monitored every day for signs of distress. The humane endpoints of the mice included decreased socialization, anorexia, weight loss of 15%, hunching, or the inability to evade handling. All hydrodynamic injections and thermal treatments were conducted under anesthesia with 5% isoflurane (Panion & BF Biotech Inc., Taipei, Taiwan), and all efforts were made to minimize suffering. For sacrifice, mice were humanely euthanized via carbon dioxide (CO_2_) gas asphyxiation followed by cervical dislocation and were considered non-survivors.

### Animals

C57BL/6 male mice (6-8 weeks old, 22-26 g) were obtained from National Laboratory Animal Center (Taipei, Taiwan). All mice were housed in groups of 6 mice per cage (32 cm × 16 cm × 14 cm) and maintained in a specific-pathogen-free (SPF) room under controlled temperature (22±2 °C) and relative humidity (55±10%) in Animal Resource Center of National Taiwan University. All mice were subjected to a 12:12 h light/dark cycle (lights on at 7:00 a.m.) with free access to food and water. All experiments were conducted with mice at age 6-8 weeks. All mice were randomly categorized into the control group, Aβ group, TC-HT group, and HT group.

### β-amyloid administration

Aβ_25-35_ (Sigma-Aldrich; Merck KGaA, Darmstadt, Germany) was dissolved in 0.9% sterile saline solution to a concentration of 1 mg/mL and then incubated at 37°C for 7 days for fibril aggregation. At the beginning of the experiment (Day 0), 10 μL Aβ solution was i.c.v.-injected into the mice’s brains of the Aβ, TC-HT, and HT groups according to the protocol described in the previous study [32]. 10 μL of sterile saline solution was i.c.v.-injected into the mice’s brains of the sham-operated control group

### HT and TC-HT treatments

To focus on the thermal effect on mice’s brain and minimize the potential interference outside of brains by the conventional whole-body hyperthermia methods [33–35], we adopted a modified method to heat mice’s heads locally. As shown in **Fig 1A**, both HT and TC-HT thermal treatments were applied by a temperature controlling and heating system equipped with a DC power supply (9303D, ABM Industries Inc., NY, USA), a temperature/process controller (PZ900, RKC instrument Inc., Tokyo, Japan), and a 19 mm × 19 mm polyimide thermofoil for heating (HK5578R4.6L12A, Minco Products Inc., Fridley, MN, USA). Mice were anesthetized with 5% isoflurane during thermal treatments. Mice’s head fur was shaved and the thermofoil was taped onto the bare skin with a thin layer of vaseline for better attachment. The thermocouple measuring tip (K-type) was located under the central region of the thermalfoil, as shown in **Fig 1B**. The actual temperatures of mice’s heads were measured via the K-type thermocouple and monitored by the temperature/process controller. For TC-HT treatment, the temperature/process controller was set to implement a 10-cycle repeated procedure, in which each cycle is consisted of a desired high-temperature stage for a period of time, followed by a natural thermal dissipating stage for 30 sec at low-temperature setting 37 °C (**Fig 1C**). In the study, the high-temperature stage in TC-HT treatment was held at 42 °C for 6 min with a 0.5 °C tolerance. For HT treatment, the temperature was kept at 42 ± 0.5 °C continuously for 1 h. Monitored temperatures of mice’s heads by HT and TC-HT thermal treatments were shown in **Fig 1D**.

**Fig 1.**
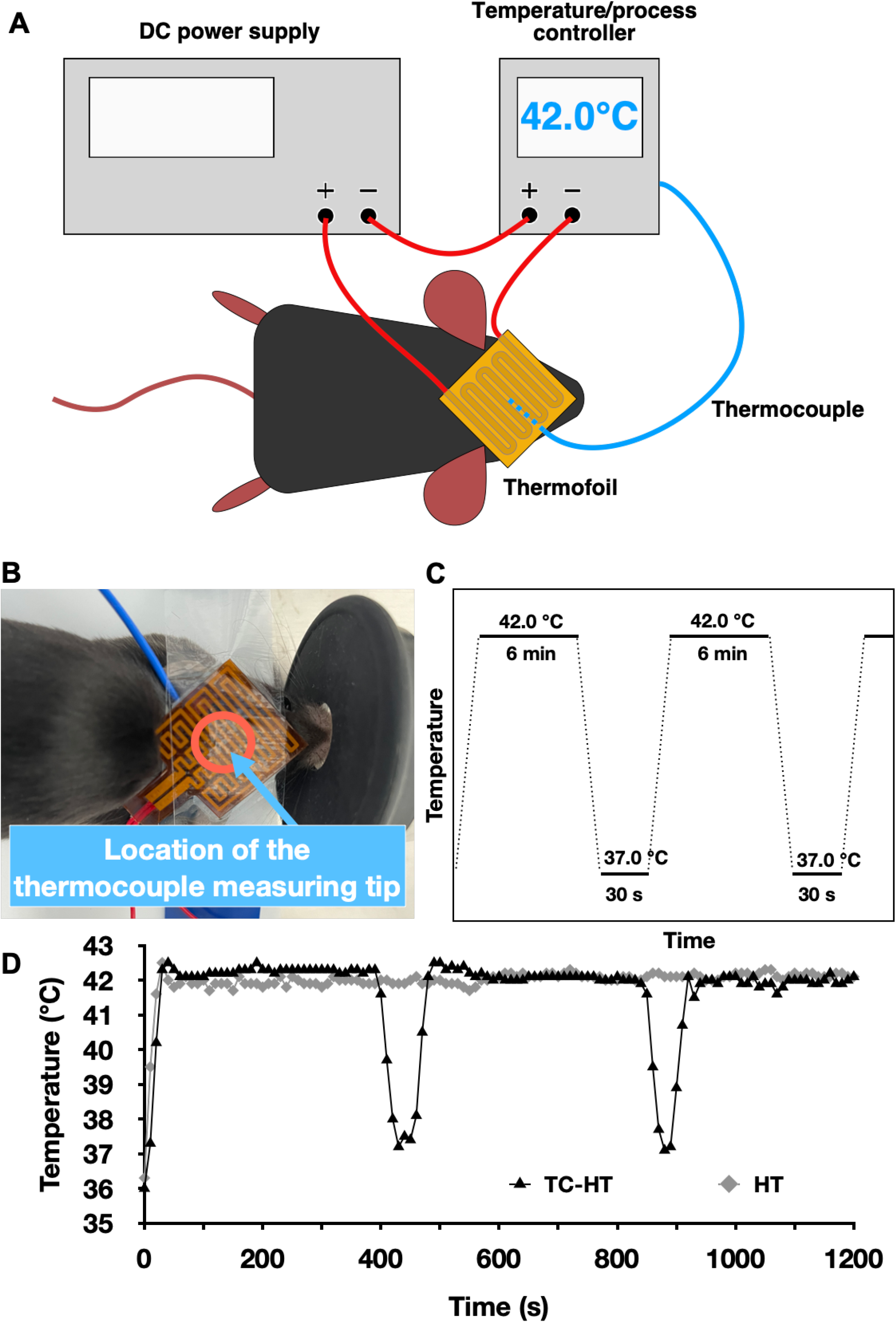
Experimental setup and design of TC-HT. (A) The setup of the temperature controlling and heating system, containing a DC power supply, a temperature/process controller, and a thermofoil for heating. (B) The thermal treatment was locally performed on mice’s heads by thermofoil, and their temperatures were measured via a K-type thermocouple, whose measuring tip was placed in the central circled region under the thermofoil. (C) Schematic diagram of the TC-HT temperature-controlled programs with parameter settings, and (D) the temperatures of mice’s heads by HT and TC-HT thermal treatments were recorded every 10 sec throughout the heating period.

### Experimental design

At the beginning of the experiment (Day 0), 10 μL Aβ solution was i.c.v.-injected into the mice’s brains of the Aβ, TC-HT, and HT groups. On day 4, 8, and 12, the respective thermal treatment was applied to the mice of the TC-HT and HT groups. Y-maze and NOR tests were assessed on day 13 and day 14, respectively. Brain tissues were collected after the Y-maze test for the following western blot analyses. The experimental schedule was shown in **Fig 2**.

**Fig 2.**
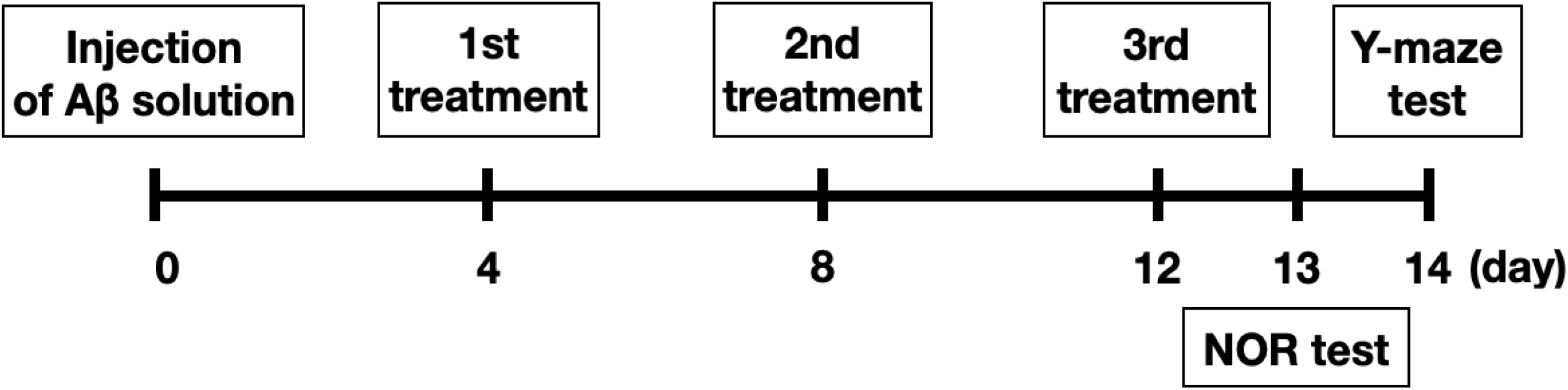
The experimental schedule. Aβ solution was i.c.v.-injected at day 0, and the respective thermal treatment was implemented at day 4, 8, and 12. Behavioral tests of NOR and Y-maze were carried out at day 13 and 14, respectively. After the Y-maze test, mice were weighted and then euthanized to collect brain tissues for the following western blot analyses.

### Y-maze test

The Y-maze test was used to assess the spatial working memory of mice. The apparatus is consisted of three identical arms (40 cm × 13 cm × 6.5 cm) interconnected at a central junction at 120° each. The natural tendency of rodents is to explore novel environments, which constitutes their spontaneous alteration behavior to explore a new arm in Y-maze. The assessed mouse was placed at the end of the start arm, and was allowed to explore all three arms freely for 8 min. Olfactory cues in the Y-maze were removed by 70% ethanol after each assessment. The behavioral performance was recorded, and the spontaneous index was calculated as follows: (number of correct alternations) / (total number of arm entries − 2) × 100. In addition, the total number of arm entries in the Y-maze was used as a measure to characterize locomotor activity.

### Novel object recognition (NOR) test

We used the NOR test to assess the recognition memory of animals. The NOR test consists of a habituation phase, a familiarization phase and a test phase. All mice in the habituation phase were placed in a 40 cm × 40 cm × 30.5 cm empty box for 8 min.

In the familiarization phase, the assessed mouse was placed in the box for 8 min with 2 identical Lego blocks, which were placed close to two diagonal corners of the box. After the familiarization phase, the mouse was returned to the home cage for 1 h and then was placed back into the testing box for the test phase. One of the original objects was replaced with a novel Lego block of a different shape in the test phase, and the behavioral performance of the mouse was recorded with an overhead video camera for 8 min. After each assessment in all phases, olfactory cues in the box and on the Lego blocks were removed by 70% ethanol. Exploration was defined as sniffing or touching the object with the nose and/or forepaws. The duration of the exploring time was calculated for the discrimination index. A discrimination index was defined as follows: (Tn-To) / (Tn+To), where Tn and To stand for the exploration time of the novel and original objects, respectively.

### Brain tissue collection and western blot sample preparation

Mice were euthanized by CO_2_ gas. After confirming the death of mice, brains were taken out quickly and rinsed with phosphate buffered saline (PBS) (HyClone; GE Healthcare Life Sciences, Chicago, IL, USA) supplemented with 1% fresh protease inhibitor cocktail (EMD Millipore, Billerica, MA, USA). Whole hippocampi were rapidly dissected on an ice-cold plate, and then stored in ice-cold RIPA lysis buffer (EMD Millipore) supplemented with 1% fresh protease inhibitor cocktail. Hippocampal tissues were homogenized via ultrasound processing (5 sec rectangular pulse with 5 sec interval for 3 times, amplitude 50%, by a Q125 sonicator; Qsonica, Newton, CT, USA) and then centrifuged at 23000×g for 30 min at 4°C. The supernatants were collected and the protein concentration was quantified by the Bradford protein assay (BioShop Canada Inc., Burlington, Ontario, Canada). Protein samples were stored at −80°C for subsequent western blotting assessments.

### Western blot analysis

Proteins from the homogenized hippocampi were resolved by sodium dodecyl sulfate polyacrylamide gel electrophoresis (SDS-PAGE) and then electroblotted onto the polyvinylidene difluoride (PVDF) membranes (EMD Millipore). Blotted membranes were blocked with 5% skim milk in Tris-buffered saline supplemented with 0.1% Tween-20 (TBST) (both from BioShop Canada Inc.) for 1 h at room temperature (RT) and then incubated overnight with diluted primary antibodies in blocking buffer at 4 °C, followed by three rinses with TBST washing buffer. Primary antibodies employed in this study were listed as following: anti-Aβ (sc-28365, Santa Cruz Biotechnology, Inc., Dallas, TX, USA), anti-IDE (ab32216, Abcam, Cambridge, UK), anti-BACE1 (#5606), anti-SOD2 (#13141), anti-Iba-1 (#17198) (Cell Signaling Technology, Danvers, MA, USA), anti-GFAP (GTX100850), and anti-β-actin (GTX110564) (Gentex, Irvine, CA, USA). After washing with TBST, the membranes were incubated with horseradish peroxidase-conjugated goat anti-mouse (for probing anti-Aβ primary antibody) (ab205719, Abcam) or goat anti-rabbit (for probing the other primary antibodies) secondary antibodies (AB_2313567, Jackson ImmunoResearch Laboratories, Inc., West Grove, PA, USA) for 1 h at RT. All the antibodies were diluted according to the manufacturer’s instructions to their optimal concentration. The protein bands were visualized by an enhanced chemiluminescence substrate (Advansta, San Jose, CA, USA) and detected by an imaging system Amersham Imager 600 (GE Healthcare Life Sciences). The images were analyzed with Image Lab software (Bio-Rad Laboratories, Inc.). For normalization of proteins, β-actin was used as the loading control to determine the relative folds of targeted proteins.

### *In vitro* cell culture and hydrogen gas (H2) treatment

The human neuroblastoma SH-SY5Y cells were purchased from American Type Culture Collection (Manassas, VA, USA). The cell culture medium was MEM/F-12 mixture (HyClone; GE Healthcare Life Sciences) containing 10% fetal bovine serum (HyClone; GE Healthcare Life Sciences) and 1% penicillin-streptomycin (Gibco Life Technologies, Grand Island, NY, USA), supplemented with 1mM sodium pyruvate (Sigma-Aldrich; Merck KGaA) and 0.1 mM non-essential amino acids (Invitrogen Life Technologies, Grand Island, NY, USA). Cells were maintained in a humidified incubator composed of 5% CO_2_ at 37°C. Before the experiment, SH-SY5Y cells were subcultured and seeded in 96-well plates (Thermo Fisher Scientific Inc., Waltham, MA, USA) overnight. Cells were treated with 25 μM Aβ for 1 h and then subjected to the TC-HT treatment for about 2 h. The TC-HT treatment was implemented by the Thermal Cycler (Applied Biosystems; Thermo Fisher Scientific, Inc.) and the *in vitro* TC-HT parameter settings were described in our previous study [31]. In the experiment, H_2_-mixed gas was composed of 75% H_2_, 20% oxygen gas (O_2_), and 5% CO_2_ gas (vol/vol/vol), as described elsewhere [36]. After continuous exposure to the H_2_-mixed gas, it has been observed that H_2_ concentration of the culture medium in a closed acrylic box gets saturated within 20~30 min at atmospheric pressure [37]. Upon 12~16 h after TC-HT treatment, cultured SH-SY5Y cells were sent into an acrylic chamber of volume about 450 mL. Mixed gas consisting of 75% H_2_, 20% O_2_, and 5% CO_2_ (vol/vol/vol) were well mixed and then flowed into the chamber at a rate of 400 mL/min for 90 min under normal pressure. Upon 4 days after the Aβ administration, the 3-(4,5-dimethylthiazol-2-yl)-2,5-diphenyltetrazolium bromide (MTT) (Sigma-Aldrich; Merck KGaA) assay was used to analyze the cell viability of SH-SY5Y cells. In brief, the culture medium was replaced with MTT solution (0.5 mg/mL in the MEM/F12 mixture) and incubated at 37°C for 4 h. The formazan crystals were dissolved by equal volume of the solubilizing buffer containing 10% sodium dodecyl sulfate (SDS) (Bioshop Canada Inc.) solution in 0.01 N hydrochloric acid (HCl) (Echo Chemical Co. Ltd., Taiwan) at 37 °C overnight. The absorbance at 570 nm of each well was analyzed by Multiskan GO microplate spectrophotometer (Thermo Fisher Scientific Inc.), and the background absorbance at 690 nm was subtracted. The cell viability of each group was expressed as percentage of the untreated control.

### Statistical analysis

All statistical results were shown as mean ± standard deviation (S.D.). Statistical analysis was conducted with OriginPro 2015 software (OriginLab Corporation, Northampton, MA, USA) using one-way analysis of variance (ANOVA), followed by Tukey’s post hoc test. *P*-value < 0.05 was considered to indicate a statistically significant difference.

## Results

### TC-HT improves Aβ-induced recognition impairments in mice

The study employed Y-maze test and NOR test to examine the therapeutic effect of TC-HT and HT on AD mice. In the Y-maze test, the overall percentage of each mouse’s spontaneous index was computed to show its short-term memory function. As exhibited in **Fig 3A**, the spontaneous index of the Aβ group is inhibited significantly, ranging from 60.3% to 44.1%, compared with the control group. TC-HT significantly improves the memory deficiency, elevating the spontaneous index to 61.6%, notably different from HT, which only causes an insignificant improvement (spontaneous index 46.8%). Besides, there is no significant difference in the total number of arm entries among the four groups (**Fig 3B**), suggesting that i.c.v. Aβ injection or both thermal treatments didn’t affect the locomotor activities of mice. On the other hand, NOR test also confirmed that TC-HT is more effective in alleviating Aβ-induced memory impairment than HT. As shown in **Fig 3C**, along with memory decline, the discrimination index of the Aβ group plunged from 25.1% down to −26.2%, which bounces back to 19.2% after TC-HT treatment, significantly outperforming HT treatment, which only raises the discrimination index to −3.1%. Thus, TC-HT exhibited a much stronger reversing effect on cognitive impairment than HT in both Y-maze and NOR tests.

**Fig 3.**
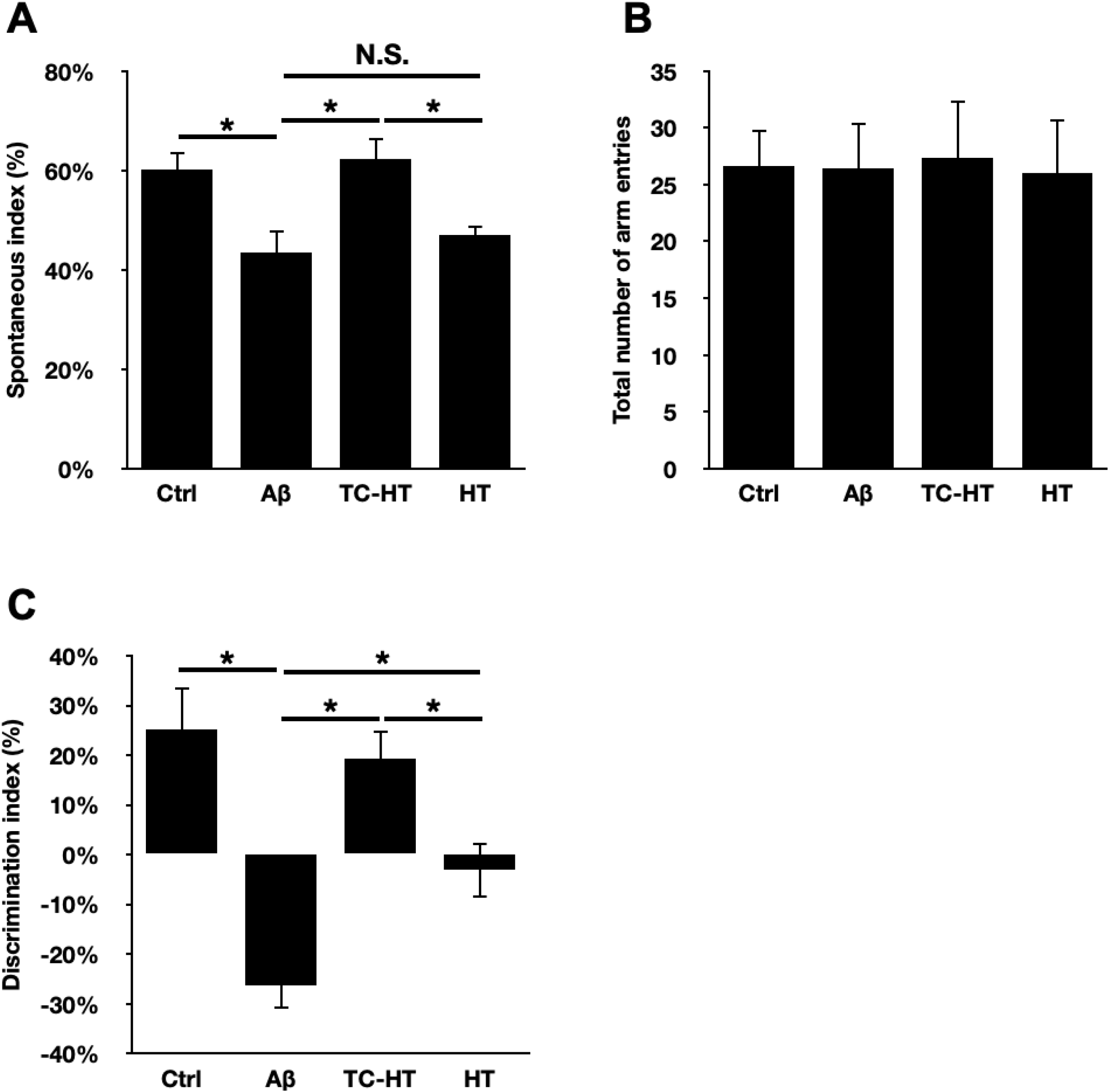
Effect of TC-HT and HT treatments on mice’s behavioral performances. The recovery effects of Aβ-induced recognition impairments after TC-HT and HT treatments are assessed via (A) the spontaneous index and (B) the total number of arm entries in the Y-maze test, and (C) the discrimination index in the NOR test. Data are presented as mean ± S.D. (n = 9 in each group). One-way ANOVA with Tukey’s post hoc test is used to determine statistical significance. (\**P* < 0.05, comparison between indicated groups).

### TC-HT down-regulates Aβ secretion and accumulation in mouse brain

The study first analyzed the accumulated Aβ levels in hippocampi, confirming the significant effect of TC-HT treatment in repairing Aβ-induced cognitive impairment, as it reduces the hippocampal Aβ level from 1.34-fold to 0.88-fold compared to the control group (**Fig 4A**). HT treatment, by comparison, exhibited only slight inhibitive effect on the hippocampal Aβ level (1.24-fold). On the other hand, it is realized that Aβ level in mouse brain is partially attributed to the pathological changes of BACE1 [38]. In **Fig 4B**, our results show that i.c.v.-injected Aβ triggers a significant increase in hippocampal BACE1 expression (1.30-fold) compared with the control group. The study finds that the elevation of BACE1 expression is significantly suppressed by the TC-HT treatment (0.89-fold), but only slightly decreased by the HT treatment (1.19-fold). It’s noteworthy that there is no significant difference in BACE1 levels between the Aβ group and the HT group, suggesting that conventional HT treatment is ineffective in stopping abnormal Aβ secretion in mouse brain. To the best of our knowledge, this is the first time that a thermal treatment exhibited an inhibitory ability on the expression level of BACE1. In sum, the study shows that TC-HT appears to be much more effective AD treatment than HT, as it’s more capable of regulating Aβ levels actively and inhibiting BACE1 level in hippocampus.

**Fig 4.**
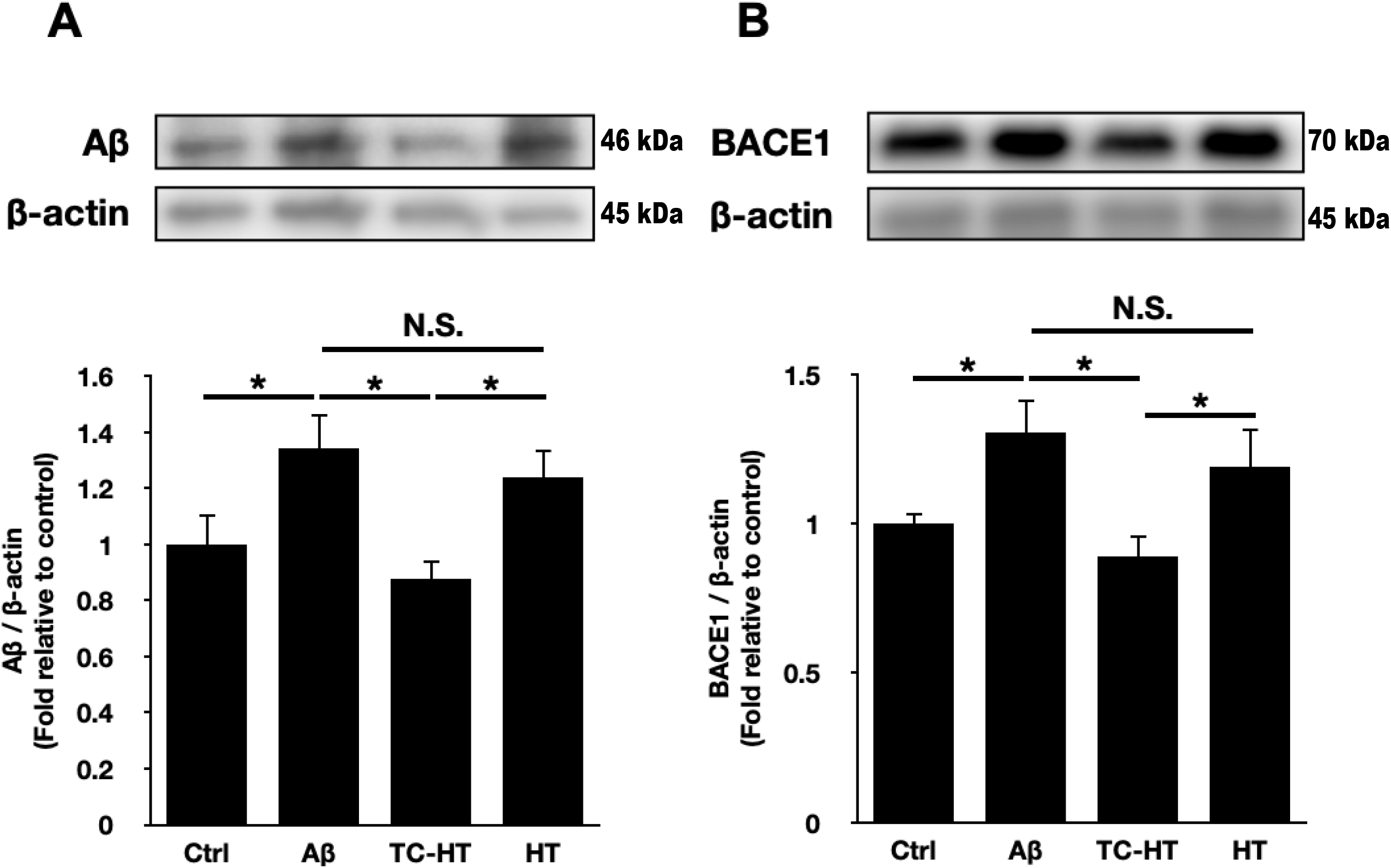
Effect of TC-HT and HT treatments on the expression levels of Aβ and BACE1 in hippocampi. Representative western blots and the quantifications of (A) Aβ and (B) BACE1 are presented for evaluating anti-Aβ effect in hippocampi. β-actin was used as loading control. Data are presented as mean ± S.D. (n = 9 in each group). One-way ANOVA with Tukey’s post hoc test is used to determine statistical significance. (\**P* < 0.05, comparison between indicated groups).

### TC-HT ameliorates Aβ-induced neuroinflammation in mouse brain

Neuroinflammation is another hallmark of AD, as the risk of cognitive impairment is highly associated with inflammatory level [39]. The study finds that the expression levels of both hippocampal Iba-1 and GFAP, the activation markers of microglia cells and astrocytes, respectively, are significantly enhanced in the Aβ group, compared with the control group (**Fig 5A** and **5B**). After the TC-HT treatment, both Iba-1 and GFAP levels drop significantly, while the HT treatment only slightly attenuates their expression levels. There is no significant difference between the Aβ group and the HT group in the expressions of both Iba-1 and GFAP. The results indicate that TC-HT treatment can better ameliorate Aβ-induced neuroinflammation than HT treatment *in vivo*.

**Fig 5.**
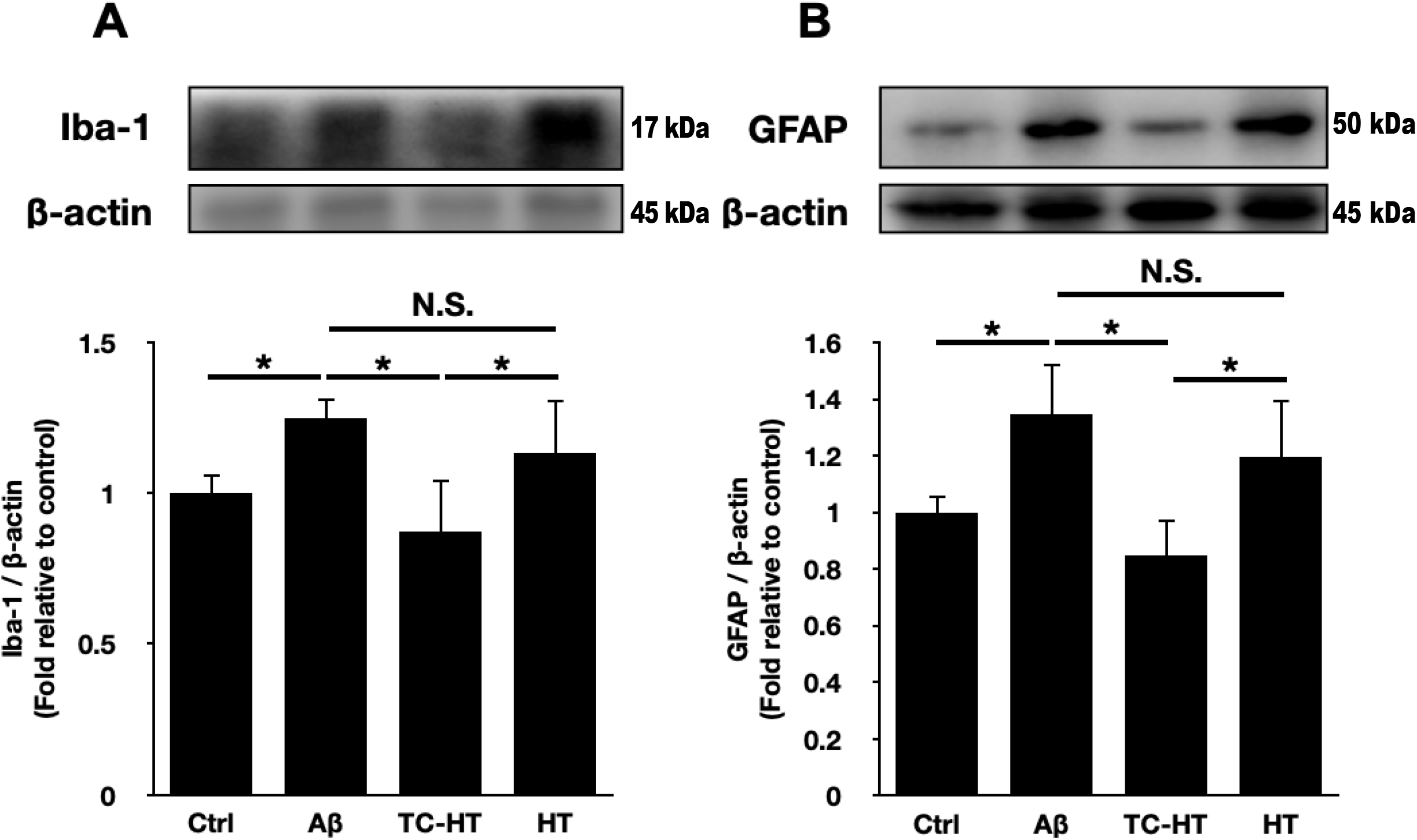
Effect of TC-HT and HT treatments on the expression levels of neuroinflammatory markers Iba-1 and GFAP in hippocampi. Representative western blots and the quantifications of (A) Iba-1 and (B) GFAP are presented for evaluating anti-neuroinflammation effect in hippocampi. β-actin was used as loading control. Data are presented as mean ± S.D. (n = 9 in each group). One-way ANOVA with Tukey’s post hoc test is used to determine statistical significance. (*\*P* < 0.05, comparison between indicated groups).

### TC-HT promotes the expressions of IDE and SOD2 proteins in mouse brain

With IDE long regarded as a major endogenous enzyme attributing to Aβ degradation, the alteration of IDE activity and expression can cause Aβ accumulation, thereby increasing the risk of AD [40]. **Fig 6A** shows that while Aβ injection slightly decreases the hippocampal IDE level (0.88-fold), TC-HT treatment significantly increases the IDE expression level in hippocampi up to 1.38-fold. By contrast, HT treatment performs an insignificant altering effect on IDE expression (0.91-fold), compared with the Aβ group. The findings show that TC-HT can intensify Aβ clearance via elevating the IDE expression level in hippocampi, while HT is ineffective in promoting the hippocampal IDE level and thus cannot contribute to the removal of abnormal hippocampal Aβ in mice.

**Fig 6.**
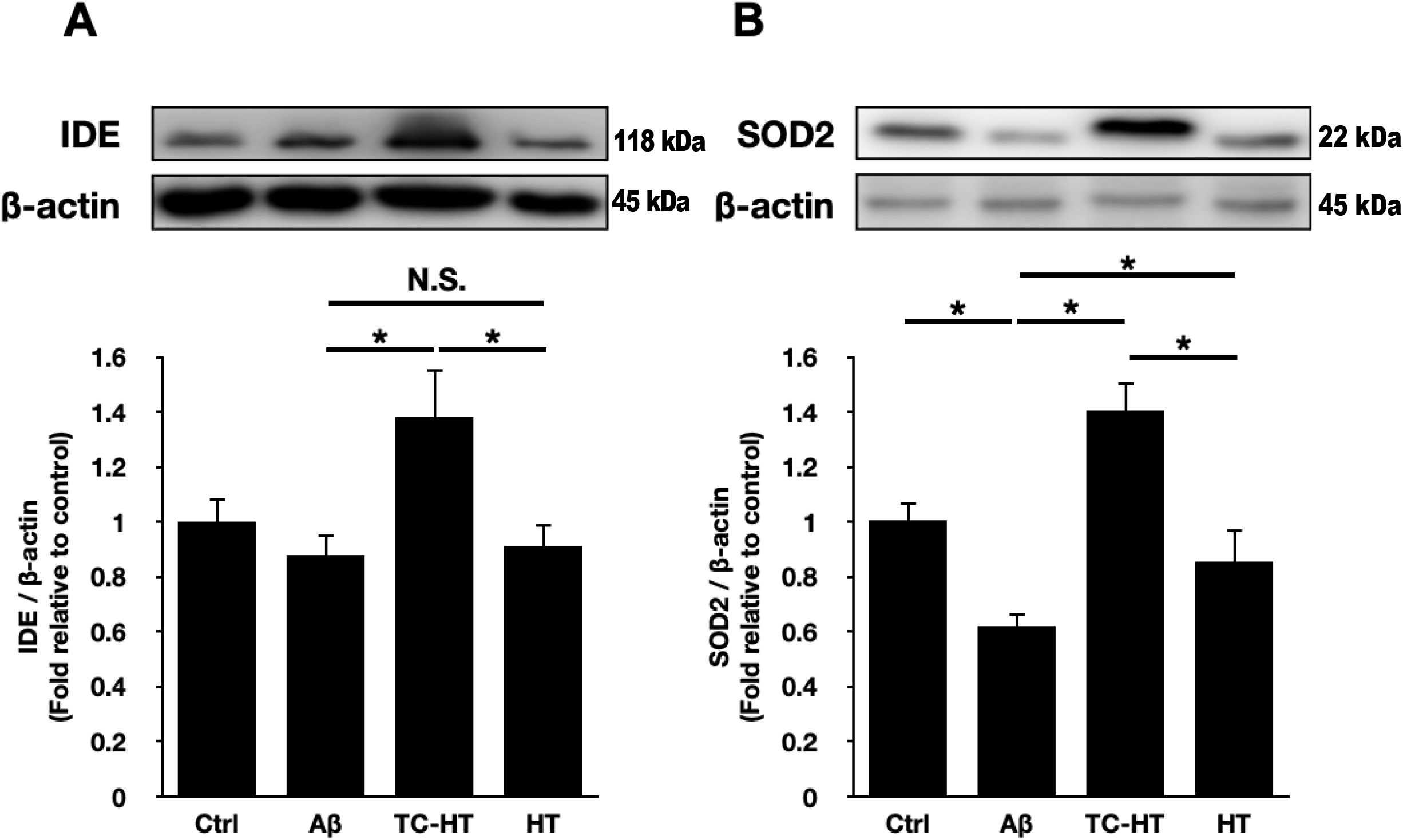
Effect of TC-HT and HT treatments on the expression levels of neuroprotective proteins IDE and SOD2 in hippocampi. Representative western blots and the quantifications of (A) IDE and (B) SOD2 are presented for evaluating Aβ clearance and antioxidant effect in hippocampi, respectively. β-actin was used as loading control. Data are presented as mean ± S.D. (n = 9 in each group). One-way ANOVA with Tukey’s post hoc test is used to determine statistical significance. (\**P* < 0.05, comparison between indicated groups).

Antioxidant protein SOD2 also has significant potential in AD treatment, capable of decreasing hippocampal superoxide and amyloid plaque in AD mice [41]. **Fig 6B** shows that after i.c.v. Aβ injection, the SOD2 expression level drops significantly (0.62-fold), which, though, recovers significantly in the TC-HT group (1.41-fold), much higher than recovery to 0.86-fold after HT treatment. The results show that TC-HT can activate stronger antioxidant activity in hippocampus against the Aβ-induced oxidative stress than continuous HT *in vivo*.

### H_2_ treatment produces auxiliary neuroprotective effect against Aβ after TC-HT treatment *in vitro*

To examine whether removal of excessive ROS after TC-HT treatment can intensify the neuroprotective effect against Aβ, H_2_ treatment as mentioned in Materials and Methods section was introduced as an exogenous ROS scavenger in an *in vitro* model. **Fig 7A** shows the significant suppression of the viability of SH-SY5Y cells to 60.9% under the insult of 25 μM Aβ, which recovers greatly to 82.5% after TC-HT treatment. Moreover, we find that with a 90 min H_2_ treatment at 12~16 h after TC-HT treatment, the cell viability can bounce back to a near-normal level (96.0%), exhibiting a better neuroprotective effect than TC-HT treatment alone. However, it should be noted that administration of H_2_ treatment less than 12 h after TC-HT treatment fails to further boost the viability of SH-SY5Y cells. On the other hand, H_2_ treatment alone does not show any significant protective effect on SH-SY5Y cells against Aβ (**Fig 7B**). These findings show that while removal of ROS via H_2_ treatment alone has no effect against Aβ, it can further enhance the neuroprotective effect of TC-HT treatment against Aβ, exhibiting a better neuroprotective effect than TC-HT treatment alone. This could be attributed to removal of TC-HT-induced ROS with H_2_ treatment after establishment of neuroprotective mechanism at 12~16 h after TC-HT treatment. These results shed light on the approach in developing AD treatment.

**Fig 7.**
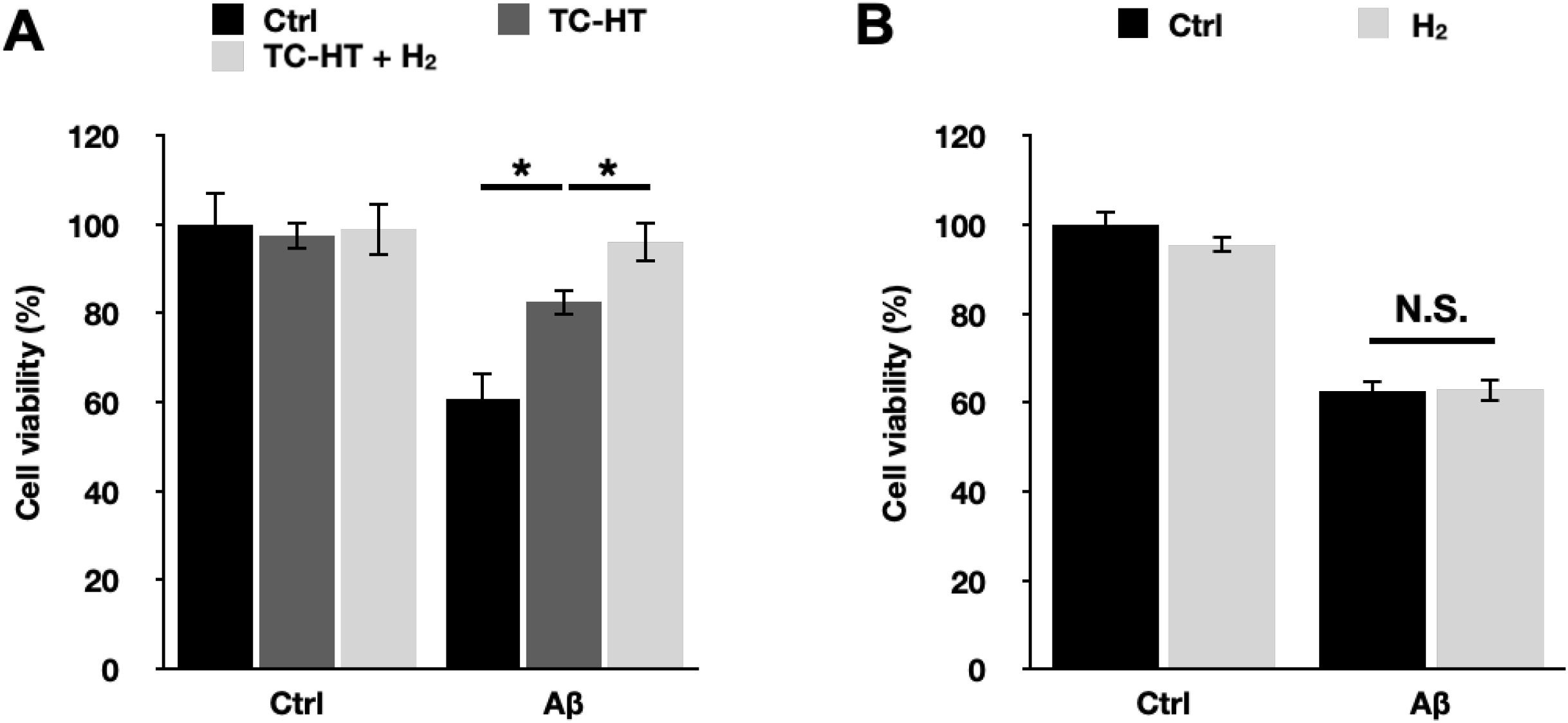
H_2_ treatment as a ROS scavenger helps to promote further the neuroprotective effect of TC-HT against Aβ *in vitro*. (A) Cell viabilities of SH-SY5Y cells treated with Aβ, Aβ + TC-HT, and Aβ + TC-HT + H_2_ treatment, respectively. The data show that H_2_-assisted TC-HT treatment exhibits a better neuroprotective effect than TC-HT treatment alone. In the other experiment, SH-SY5Y cells were subjected directly to H_2_ treatment after the administration of 25 μM Aβ. Cell viabilities of SH-SY5Y cells treated with Aβ and Aβ + H_2_ treatment alone are respectively shown in (B). The result indicates that H_2_ treatment alone has no effect against Aβ. Data are presented as mean ± S.D. (n = 6 in each group). One-way ANOVA with Tukey’s post hoc test is used to determine statistical significance. (\**P* < 0.05, comparison between indicated groups).

## Discussion

Given prevalence of AD in our society in recent years, many scientists have taken part in research on its treatment with various approaches, including Aβ hypothesis, one of the most noticeable pathogenesis of AD [42]. Aβ peptides could be secreted in brains, but their abnormal depositions, especially in hippocampus, could induce severe neurodegeneration and synapse loss, thereby impairing memory and cognitive abilities [7, 8]. The study employed Aβ_25-35_, peptide fragment of Aβ_1-42_ considered the most pathogenic Aβ species in AD [43, 44], as the model. Development of AD drugs has been confronted with two major obstacles, namely blood-brain barrier (BBB) and severe side effects [14, 45]. Hence, the application of heat has potential as an alternative AD treatment, due to its capability to intensify neuroprotective effect [31] and sidestep BBB, targeting the diseased brain directly. The study embraced TC-HT, a modified heat treatment with higher safety and superior protective effect than HT [31], to investigate the therapeutic potential of thermal treatment on AD *in vivo*.

With memory loss being the first and main symptom of AD [46], AD studies with mice as model have often employed Y-maze and NOR tests to assess short-term memory and learning memory [47, 48]. The study probed the therapeutic effect of HT and TC-HT treatment on Aβ-induced memory impairment, finding that TC-HT treatment can significantly improve the performance of spontaneous index (**Fig 3A**) and help memory-impaired mice recognize and remember new objects (**Fig 3C**). The study demonstrates significant effect of TC-HT treatment in recovering spatial working memory and recognition memory impaired by Aβ. In comparison, although HT treatment also enhances behavioral performance, the extent of improvement significantly lags behind that of TC-HT treatment in both tests. Collectively, TC-HT treatment exhibits a higher therapeutic potential than HT treatment in improving Aβ-induced memory deficiency and cognitive dysfunction *in vivo*.

From a molecular perspective, Aβ hypothesis has long been a mainstream approach in the study on the mechanism of AD pathogenesis and drug development [49]. Under normal conditions, α- and γ-secretase will hydrolyze APP, preventing production of Aβ peptides. However, once pathological changes occur, the hydrolysis of APP is carried out by β- and γ-secretase, leading to secretion of toxic Aβ peptides and aggregation of insoluble Aβ fibrils [3]. Thus, under the Aβ hypothesis, inhibition of BACE1 is also considered a major therapeutic target, on top of Aβ clearance, in AD treatment. The study exhibits the significant effect of TC-HT in decreasing the hippocampal Aβ level, remarkably outperforming HT which fails to show any significant Aβ clearance ability (**Fig 4A**). Besides, TC-HT also shows a significantly better effect in inhibiting the hippocampal BACE1 level than HT, and there is no significant difference in BACE1 level between the HT and Aβ groups (**Fig 4B**). These results demonstrate that TC-HT can attenuate the amount of toxic Aβ fragments and their production in the brains of AD mice, much better than the continuous HT treatment. To the best of our knowledge, this is the first time that a physical thermal treatment is proven to be able to modulate the expression of BACE1, shedding light on alternative approach in AD treatment.

Apart from neurotoxic Aβ in brain, the pathogenesis of AD is also closely linked to the activation of the brain’s immune cells, particularly microglia and astrocytes, which are triggered by Aβ deposits, according to some reports [4, 50]. Chronic activation of the immune cells causes neuroinflammation, releasing the inflammatory mediators which are associated with neurodegeneration and thus exacerbating AD progression [50, 51]. Consequently, inhibition of neuroinflammation in AD brains is also important in AD treatment. The study demonstrates extraordinary anti-neuroinflammatory ability of TC-HT treatment. It is capable of suppressing both Aβ-induced activation markers of microglial cells (Iba-1) and astrocytes (GFAP) significantly, as shown in **Fig 5**. By contrast, the continuous HT treatment shows only little inhibition on both neuroinflammatory markers without significance. Therefore, the study proves that TC-HT is more capable of ameliorating neuroinflammation in AD brains than continuous HT *in vivo*.

The clearance of toxic Aβ deposits, a hallmark of Alzheimer’s disease, is as important as curbing its production. First discovered as an enzyme responsible for the degradation of insulin, IDE has also been found in recent studies to be capable of degrading several polypeptides including Aβ [40]. In addition, it has been known that neuron cells are able to regulate the extracellular levels of Aβ via proteolysis by IDE *in vitro* [52]. Studies have also shown that the alteration of IDE plays a key role in AD, as the decrease of IDE activity and expression could result in Aβ accumulation, thereby raising the risk of AD development [53, 54]. In line with the finding of a recent study that IDE can be increased in a fashion like HSP [21], our study discovers that TC-HT treatment can significantly boost the hippocampal IDE level (**Fig 6A**), an effect unfound in HT treatment. The more pronounced Aβ clearance effect of TC-HT may result from its higher capability in inhibiting Aβ production (**Fig 4B**) and clearing Aβ (**Fig 6A**) than continuous HT, indicating its higher potential in AD therapy than the latter.

Antioxidant enzyme is also an important protective mechanism against AD. It has been known that Aβ deposition in brain can increase the production of mitochondrial ROS and intracellular oxidative stress, increasing Aβ formation, which leads to a vicious cycle and exacerbates AD progression [5, 6]. Besides, excessive ROS level could be toxic to brain neurons and decrease cognitive performance, according to some studies [55]. SOD2, a member of the superoxide dismutase family, can transform superoxide O_2_ · ^-^ into hydrogen peroxide and hence regulate excessive oxidative stress [56]. The reduction of SOD2 can accelerate the onset of Aβ-dependent memory deficits *in vivo* [57], while the elevation of SOD2 can prevent the memory deficits [58]. Our study shows that TC-HT can greatly recover hippocampal SOD2 decreased by Aβ (**Fig 6B**), at a scale far outpacing that of HT. The result is in line with the behavioral performance of mice in Y-maze and NOR tests in **Fig 3**, indicating that TC-HT can attenuate the Aβ-induced memory deficits via removal of excessive ROS in brains by SOD2-related mechanisms.

In addition to the neuroprotective functions of IDE and SOD2, silent mating-type information regulation 2 homolog 1 (SIRT1) has been reported to attenuate Aβ levels both *in vitro* and *in vivo* [59]. SIRT1 is a famous deacetylase, known for removing acetyl groups from substrate proteins to regulate their function. The SIRT1-mediated deacetylation can not only reduce Aβ production by suppressing the expression of β-secretase, but also inhibit tau protein aggregation, another pathological hallmark of AD [59, 60]. Besides, SIRT1 is capable of suppressing inflammation and activating antioxidative response via deacetylation [61, 62]. Noteworthily, the deacetylation activity of SIRT1 can be induced by heat [63, 64], which could deacetylate heat shock factor 1 and lead to gene transcription of heat shock proteins such as HSP70 [64].

HSP70 is a well-known heat-induced chaperone protein that assists proper protein folding or refold the misfolded proteins in cells [65]. In addition to its role in protein folding, HSP70 is also capable of blocking the oligomerization of Aβ fibril via refolding, clearing Aβ through up-regulating IDE expression, inhibiting tau protein aggregation, and promoting tau protein degradation [66]. Hence, we believe that both SIRT1 and HSP70 are also involved in the therapeutic effects of TC-HT against AD. Our results show that i.c.v. Aβ injection significantly reduces hippocampal SIRT1 level to 0.76-fold of the control group, which is significantly recovered by TC-HT treatment (0.96-fold) (**Fig 8A**). On the other hand, HSP70 expression is not affected in mice hippocampi by Aβ injection (1.08-fold), but it is significantly elevated in the TC-HT group (1.32-fold), as shown in **Fig 8B**. Noteworthily, the increase in HSP70 level by TC-HT treatment is consistent with the change of IDE level in **Fig 6A**, suggesting that TC-HT-induced Aβ clearance might be mediated via HSP70-dependent signalling.

**Fig 8.**
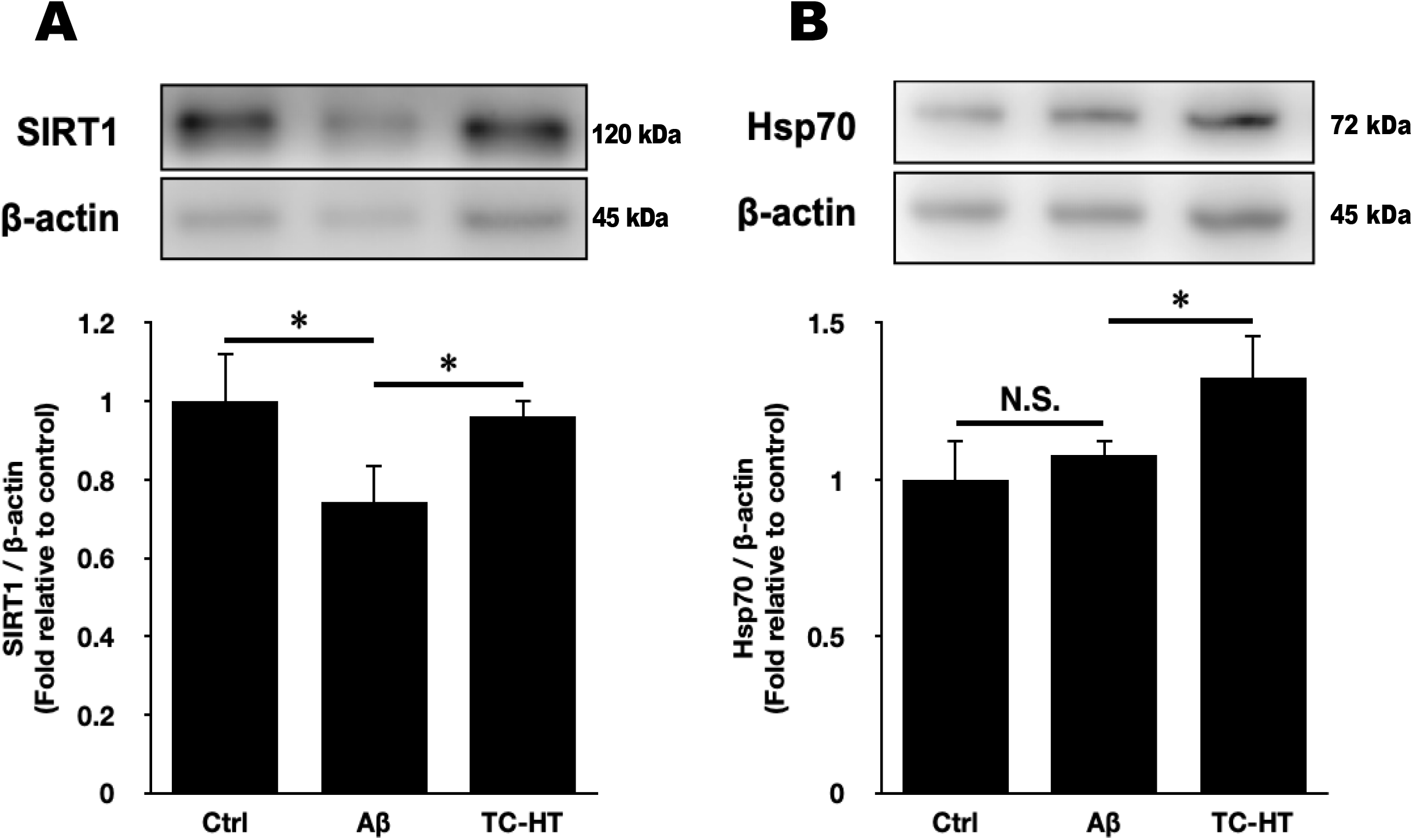
Effect of TC-HT treatments on the expression levels of SIRT1 and HSP70 in hippocampi. Representative western blots and the quantifications of (A) SIRT1 and (B) HSP70 are presented for evaluating anti-Aβ effect in hippocampi. β-actin was used as loading control. Data are presented as mean ± S.D. (n = 9 in each group). One-way ANOVA with Tukey’s post hoc test is used to determine statistical significance. (\**P* < 0.05, comparison between indicated groups).

Excessive ROS accumulation could be detrimental to neurons, damaging memory and cognitive ability. However, some studies have shown that proper increase of ROS level is essential for activating the redox signalling and the associated neuroprotective pathways [67]. Therefore, the study suggests not to eliminate ROS induced by thermal stress, until setup of relevant protective mechanisms. To validate the hypothesis, an *in vitro* experiment using H_2_ treatment as an ROS scavenger was introduced in this study. As shown in **Fig 7A**, the result shows significant recovery of the Aβ-inhibited viability of SH-SY5Y cells after TC-HT treatment, whose neuroprotective effect could be attributed to activation of Akt and its downstream signalling pathways, as suggested in our previous study [31]. It has been reported that early administration of antioxidants may block the construction of neuroprotection via these pathways [68–70], which is in line with the observation that administration of H_2_ treatment less than 12 h after TC-HT fails to further augment the viability of SH-SY5Y cells. On the other hand, our study shows that with 90 min H_2_ treatment at 12~16 h after TC-HT, removal of TC-HT-induced ROS can further boost protection of neural cells against Aβ, exhibiting a better neuroprotective effect than TC-HT treatment alone (**Fig 7A**). Noticeably, the H_2_ treatment alone did not show any protective effect for SH-SY5Y cells against 25 μM Aβ (**Fig 7B**), indicating that TC-HT treatment is essential for the protective mechanism. The results show that scavenging TC-HT-induced ROS with H_2_ treatment after establishment of neuroprotective mechanism by TC-HT (12~16 h after TC-HT treatment) can further enhance the neuroprotective effect of TC-HT treatment against Aβ. Relevant studies have been in progress *in vivo*.

Although heat could be beneficial for AD, continuous heat exposure may hinder the function of neuroprotective proteins. Some studies have shown that the inactivation of proteins may start around 40°C, due to protein denaturation and unfolding [71]. In the study, continuous HT treatment doesn’t exhibit significant effect in Aβ regulation (**Fig 4 & Fig 6A**) and anti-neuroinflammation activity (**Fig 5**), and lags behind TC-HT treatment in antioxidative activity (**Fig 6B**). However, with identical heat exposure time, TC-HT featuring intermittent high-temperature treatment attains much greater improvement than continuous HT. As a result, despite its contribution to brain’s neuroprotective mechanism, HT treatment may entail overheating causing protein dysfunction in neural cells and thus interrupting the pathways for neuroprotection. Therefore, the interspersing low-temperature process is crucial for TC-HT treatment’s neuroprotective effect. Our study demonstrates that TC-HT can be an effective thermal treatment for AD. In practice, focused ultrasound (FUS) may be a suitable heating source for TC-HT treatment, as it is capable of precisely modulating the temperature at the targeted area, via computer-controlled programmable procedure. The FUS heating area can be set at 4-8 mm [72, 73], smaller than human brain, thereby quickening heat dissipation to surrounding tissues and preventing hyperthermic damage. In operation, multi-focal FUS can control the heating area and temperature by changing the size of scanned spot and intensity in scanning mode, thereby attaining set temperature and effective thermal dissipation for TC-HT in AD treatment.

In summary, our study puts forth a novel hyperthermic method, TC-HT, for application as a potential AD treatment *in vivo*, which demonstrates a better effect in improving the cognitive performance of mice in Y-maze and NOR tests than conventional HT treatment, which heats the subjects continuously. Furthermore, the study shows that TC-HT can suppress the hippocampal Aβ expression to a normal level, significantly outperforming HT which only exhibits a slight inhibition effect, perhaps due to TC-HT’s higher capability in inhibiting the Aβ productive activity of BACE1 protein and promoting the Aβ clearance effect of IDE protein. In addition, TC-HT can significantly inhibit the Aβ-induced overexpression of neuroinflammatory markers Iba-1 and GFAP in hippocampi, while HT treatment only slightly attenuates their expression levels without significance. Moreover, TC-HT raises the expression level of antioxidative protein SOD2 at an extent much higher than HT, indicating that it can provide a better protection for hippocampal neurons against the i.c.v.-injected Aβ. Overall, the study validates the therapeutic potential of TC-HT for AD *in vivo*, shedding light on development of alternative AD treatment with high efficacy.

## Ethics Approval

This study was approved by the Institutional Animal Care and Use Committee of National Taiwan University. The experiments were conducted with Guide for the Care and Use of Laboratory Animals being strictly followed.

## Acknowledgments

The authors would like to acknowledge the service provided by Animal Resource Center of National Taiwan University for facility support.

## Author Contributions

**Conceptualization:** Chih-Yu Chao.

**Data Curation:** Yu-Yi Kuo, Chih-Yu Chao.

**Formal analysis:** Yu-Yi Kuo, Wei-Ting Chen, Guan-Bo Lin, You-Ming Chen, Hsu-Hsiang Liu, Chih-Yu Chao.

**Funding acquisition:** Chih-Yu Chao.

**Investigation:** Yu-Yi Kuo, Wei-Ting Chen, Guan-Bo Lin, Chih-Yu Chao.

**Project Administration:** Chih-Yu Chao.

**Supervision:** Chih-Yu Chao.

**Validation:** Yu-Yi Kuo, Wei-Ting Chen, Guan-Bo Lin, You-Ming Chen, Hsu-Hsiang Liu

**Writing – original draft:** Yu-Yi Kuo, Chih-Yu Chao.

**Writing – review & editing:** Chih-Yu Chao.

